# Alternate primers for whole-genome SARS-CoV-2 sequencing

**DOI:** 10.1101/2020.10.12.335513

**Authors:** Matthew Cotten, Dan Lule Bugembe, Pontiano Kaleebu, My V.T. Phan

**Affiliations:** UK Medical Research Council–Uganda Virus Research Institute and London School of Hygiene and Tropical Medicine Uganda Research Unit, Entebbe, Uganda; Uganda Virus Research Institute, Entebbe; Erasmus Medical Center, Rotterdam, the Netherlands; UK Medical Research Council–University of Glasgow Centre for Virus Research, Glasgow, Scotland, UK

**Keywords:** SARS-CoV-2, COVID-19, primers, next generation sequencing

## Abstract

As the world is struggling to control the novel Severe Acute Respiratory Syndrome Coronavirus 2 (SARS-CoV-2), there is an urgency to develop effective control measures. Essential information is encoded in the virus genome sequence with accurate and complete SARS-CoV-2 sequences essential for tracking the movement and evolution of the virus and for guiding efforts to develop vaccines and antiviral drugs. While there is unprecedented SARS-CoV-2 sequencing efforts globally, approximately 19 to 43% of the genomes generated monthly are gapped, reducing their information content. The current study documents the genome gap frequencies and their positions in the currently available data and provides an alternative primer set and a sequencing scheme to help improve the quality and coverage of the genomes.

## Introduction

Since the first report on 30^th^ December 2019 in Wuhan China and the WHO declaration of the pandemic on 12^th^ March 2020, the novel Severe Acute Respiratory Syndrome Coronavirus 2 (SARS-CoV-2) (1) and the associated disease Coronavirus Disease 2019 (COVID-19) (2)(3) have continued to spread throughout the world, causing >46 million infections and >1,200,000 death globally (4). The virus genome sequences carry important information, which can be used to interpret the virus transmission, evolution patterns and origin tracing. Furthermore, accurate and complete genomic sequences are essential for monitoring diagnostics and developing novel therapeutics and vaccines. We have seen an unprecedented amount of virus sequencing with over 130,000 complete or nearly complete genome sequences of SARS-CoV-2 now available in the GISAID database by the end of September 2020 (5). Most of the sequences have been generated by next generation sequencing using targeted amplicon methods. A scan through SARS-CoV-2 genomes from GISAID with the filter “complete genome” revealed a high frequency of gaps occurring across the genome, influencing the overall genomics quality and interpretation. Here we describe an alternate primer scheme for whole-genome sequencing to improve the genome sequence quality and coverage.

## Documenting the problem

We retrieved genomes deposited to GISAID in September 2020 (9 months into the pandemic), using the “complete genome” filter and sorting the genomes by sequencing platforms information included in the metadata. Figure 1 illustrated the positions across the 30kb genome of every stretch of 200 Ns (N200; nearly the size of an amplicon) in the first genomes deposited in September 2020 using the Illumina platform (Panel A; N = 3000) versus MinION platform (Panel B; N = 3000). Histograms of gap frequencies across the genomes are shown for each platform. The gaps are not randomly distributed but occur with higher frequency in a subset of positions across the genome. Although genomes generated by the two platforms (Illumina and MinION) show similar problem regions (nt 8000-11000, and nt 19,000-24,000 relative to the reference genome NC_045512), the patterns are not completely identical. Given the use of several primer amplification schemes, we suspect the gaps in coverage may be due to unexpected primer interactions, complicated sequence regions (odd composition or secondary structure), issues with primer trimming during quality control of read data, or some combinations of these factors.

**Figure 1.**
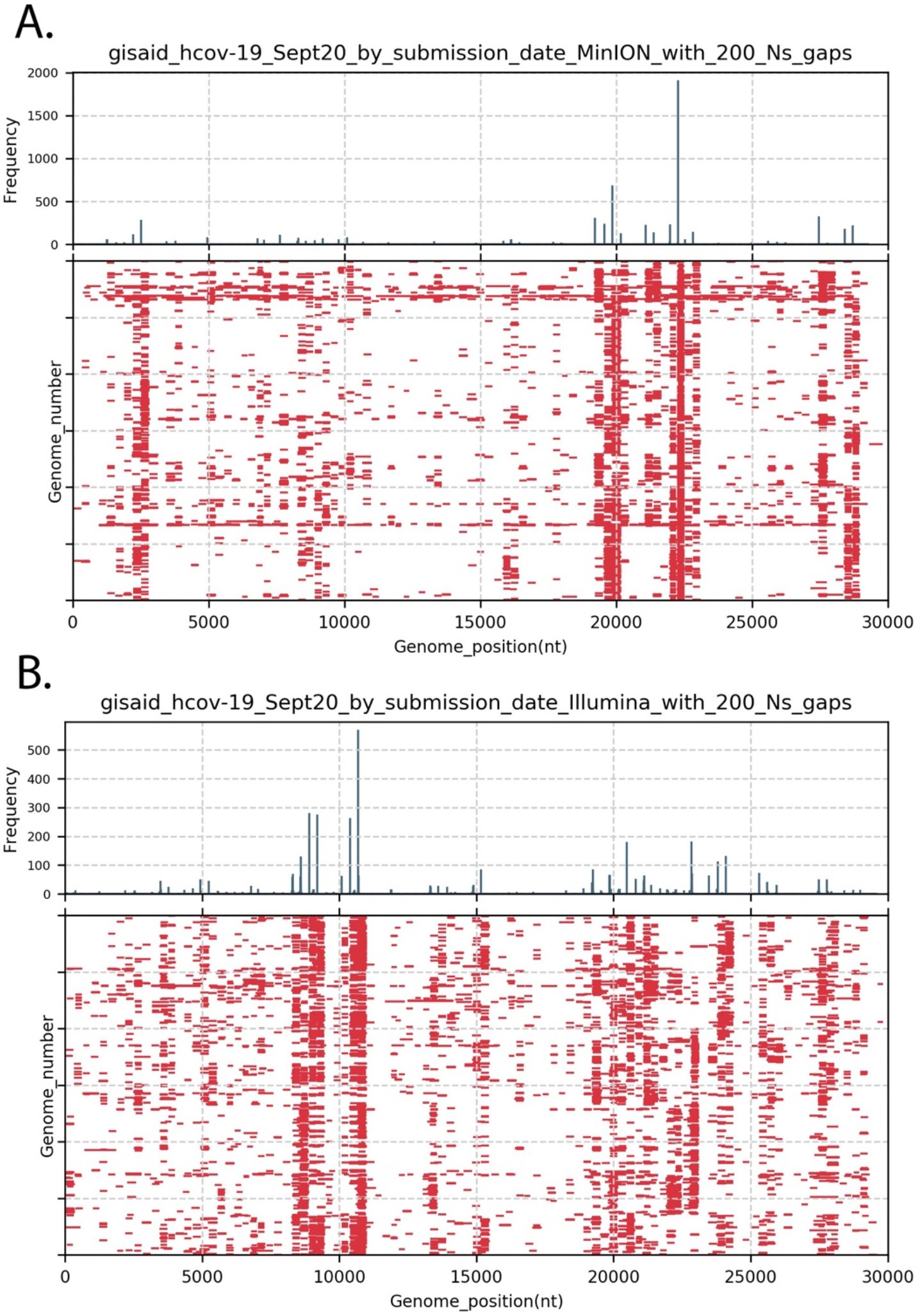
Positions of 200nt gaps across SARS-CoV-2 genomes listed as complete in GISAID. Genomes deposited in September 2020 (n= 38,228) were retrieved from GISAID, sorted by sequencing platform (MinION versus Illumina) and genomes with at least 1 instance of 200N were collected. Panel A present gaps in the first 3000 MinION-generated genome sequences deposited that contained at least on 200N motif. Gaps >=200nt in each genome are indicated with red bars. The upper panel histogram shows the frequency (in 30 nt bins) of gaps >=200nt motifs by start position on genomes. Panel B is the same analysis of the first 3000 Illumina-generated genome sequences in September 2020 that contained at least one 200N motif.

The phenomenon is unlikely to be due to an isolated set of genomes as we observed similar N200 frequencies in genomes submitted from each month of the pandemic (Table 1), suggesting that gaps in coverage is a more general phenomenon. Of note, genomes generated using Ion Torrent show much lower levels of N200 (Table 1). The very low frequency of large gaps in the Ion Torrent data may be due to the use of a dedicated alternative primer set (6). There have been discussions and reports on the SARS-CoV-2 genome changes due to sequencing errors as well as long gaps in the genomes due to missing amplicons from the amplicon-approach sequencing (7) (8). Updates of the ARTIC primers have been presented in late March 2020 to address these issues (9) (10). Additional reports of longer amplicon methods have been published (11) (12) (14) including methods to use a subset of ARTIC primers to generate longer amplicons(13).

**Table 1.** Frequency of SARS-CoV-2 genomes with 1 or more 200 nt gaps (N200) by month and by sequencing platform.

| Deposition period | Complete genomes | Genomes with 1 or more N200<sup>2</sup> | % with 1 or more N200<sup>3</sup> | Illumina total<sup>4</sup> | Illumina % with N200<sup>5</sup> | MinION total<sup>6</sup> | MinION % with N200<sup>7</sup> | Ion Torrent total<sup>8</sup> | Ion Torrent % with N200<sup>9</sup> | Method unclear<sup>10</sup> | Method unclear % with N200<sup>11</sup> |
| --- | --- | --- | --- | --- | --- | --- | --- | --- | --- | --- | --- |
| 1Jan20-31Jan20 | 54 | 1 | 2 | 9 | 0 | 2 | 0 | 0 | 0 | 43 | 2 |
| 1Feb20-29Feb20 | 126 | 2 | 2 | 44 | 0 | 9 | 0 | 5 | 0 | 65 | 4 |
| 1Mar20-31Mar20 | 2872 | 559 | 19 | 1518 | 16 | 548 | 32 | 35 | 6 | 771 | 18 |
| 1Apr20-30Apr20 | 12411 | 3745 | 30 | 4970 | 38 | 1286 | 27 | 264 | 0 | 5424 | 26 |
| 1May20-31May20 | 19787 | 8606 | 43 | 8634 | 52 | 2634 | 30 | 529 | 0 | 7990 | 42 |
| 1Jun20-30Jun20 | 21665 | 8723 | 40 | 7043 | 36 | 3844 | 35 | 629 | 2 | 10149 | 47 |
| 1Jul20-31Jul20 | 17986 | 4834 | 27 | 4965 | 23 | 1585 | 33 | 471 | 2 | 10965 | 29 |
| 1Aug20-31Aug20 | 17276 | 4005 | 23 | 11074 | 22 | 2270 | 26 | 486 | 0 | 3446 | 28 |
| 1Sep20-30Sep20 | 38227 | 10611 | 28 | 22740 | 23 | 7973 | 44 | 580 | 1 | 6934 | 28 |

1. Number of genomes with the annotation "complete" retrieved from GISAID (https://www.gisaid.org/). 2. Genomes were sorted by the presence or absence of the sequence N200. 3. $((\text{The number of genomes with at least 1 N200}) / \text{total number of genomes}) * 100$ . 4. Number of genomes in GISAID for this period generated using any of the Illumina methods as noted in the GISAID "Sequencing technology metadata" . 5. $((\text{The number of Illumina genomes with at least 1 N200}) / \text{total number of genomes}) * 100$ . - 6.,7. Number of genomes in GISAID for this period generated using any of the MinION methods as noted in the GISAID "Sequencing technology metadata" and their percentage. - 8.,9. Number of genomes in GISAID for this period generated using any of the Ion Torrent methods as noted in the GISAID "Sequencing technology metadata" and their percentage. - 10.,11. Number of genomes in GISAID for this period generated using unclear methods as noted in the GISAID "Sequencing technology metadata" and their percentage.

However, the percentage of reported complete genomes in GISAID with 1 or more N200s continues, with 10,611 (28%) of the 38,228 genomes deposited in September 2020 having 1 or more N200 gaps (Table 1), indicating the challenges remain largely unsolved.

## Detailed analysis of gaps

A more focused analysis of the frequent gaps is provided in Figure 2. The gap pattern between nt 19,000 and 24,000 (relative to the reference genome NC_045512) is shown for both MinION and Illumina sequences (first 3000 genomes of each deposited in September 2020 with at least 1 200N motif). For reference the positions of the ARTIC primers (v.1) in the region are indicated (middle panel). A histogram of gap start positions (top panel) and the individual genome gaps (bottom panel) are also shown.

**Figure 2.**
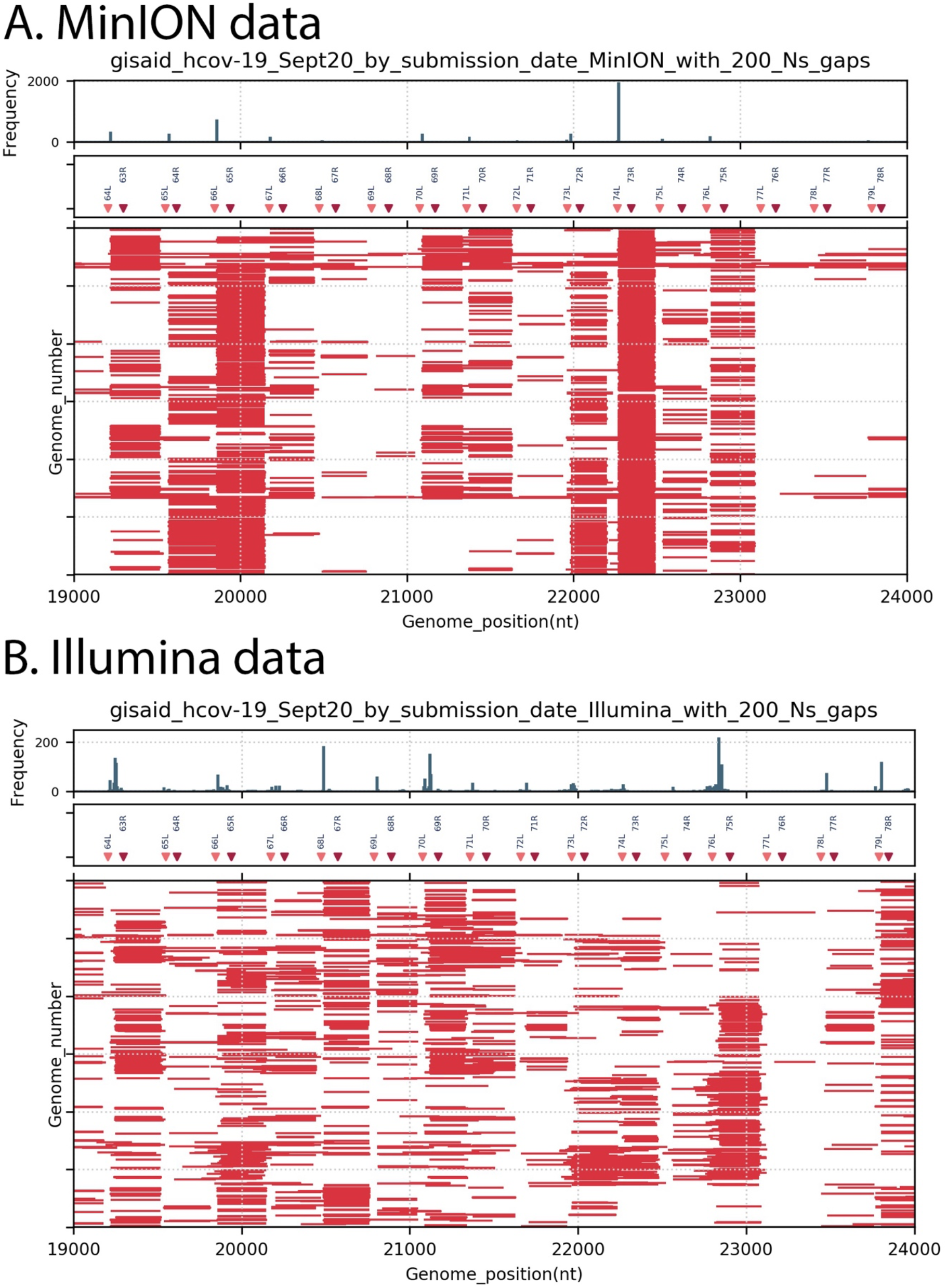
Positions of 200nt gaps across SARS-CoV-2 genomes stratified by MinION or Illumina, in region nt 19000 to 24000. Genomes deposited in September 2020 as “complete” were retrieved from GISAID, sorted by sequencing platform and by the presence of at least one N200 motif. For clarity only the first 3000 genomes in each set were plotted. Similar to Figure 1, gaps >=200nt in each genome are indicated with red bars. The upper panel histogram shows the frequency (in 30 nt bins) of gaps >=200nt motifs by start position on genome, the middle panel plots the positions of ARTIC v.1 primers in the region(pink = forward “left” primers, red= reverse “right” primers). Panel A: MinION-derived genome sequences, Panel B: Illumina-derived genome sequences.

The peaks of gap start positions frequently lie between forward primerL from Amplicon n and reverse primerR from Amplicon n-1, for both MinION and Illumina data. Because of overlapping amplicons commonly used, if a single amplicon is missing from the sequencing library (amplicon 74 for example), the resulting gap in coverage would not be the complete amplicon 74 but would span from the 3’ end of the adjacent amplicon 73 (after primer and quality trimming) to the 5’ end of adjacent amplicon 75 (after primer and quality trimming). The calculated gaps generated by such amplicon loss have a median length of 270.5 nt, which is close to the the observed median gap length in the MinION data (258 nt) or Illumina data (262 nt) from September 2020. This arrangement is outlined in the Supplementary Material Figure 1, Panel A.

## Alternate primers as a potential solution to avoid gapped genomes

We explored an alternate set of amplification primers (termed the **Entebbe primers**) designed using methods we had previously used for MERS-CoV (15), Norovirus (16), RSV (17) and Yellow Fever virus (18). Important for the design were the amplicon size and the primer placement. For their implementation, the use of primers for the reverse transcription step, and the multiplexing of the amplicons in two staggered sets was important for the PCR. Our experience had suggested an optimum amplicon size of around 1500 bp. The larger amplicons reduced the total primers content of the reactions but still allowed high reverse transcription efficiency (which, in our hands, declined beyond 1500 nt). Here we describe primers designed for whole-genome sequencing of SARS-CoV-2, as well as sharing the detailed laboratory methods that we used for reverse transcription, PCR amplification and MinION library preparation to successfully sequence the SARS-CoV-2 genome.

Briefly, the primer design (Figure 3A) started with the set of complete SARS-CoV-2 genome sequences available in the GISAID database on 22 June 2020 (N = 21,687). Spaces and disruptive characters were removed from the sequence IDs and the sequences were further screened to remove genomes containing gaps of 6Ns or more, resulting in 17,220 clean genome sequences. Next, all sequences were sliced into 33nt strings (33mers), with a 1nt step and 606,389 unique 33mers were generated. The frequency of each 33mer was counted to identify highly conserved 33mers. This counting method avoids the multiple sequence alignment step commonly used in primer design and becomes prohibitive with large and or diverse genome sets. This alignment-free approach allowed us to use all suitable genome sequences of interest rather than a set that could be conveniently aligned. Finally, primer-like 33mers sequences were generated by trimming the sequences to a calculated desired melting temperature and removing any primers greater than 26nt.

**Figure 3.**
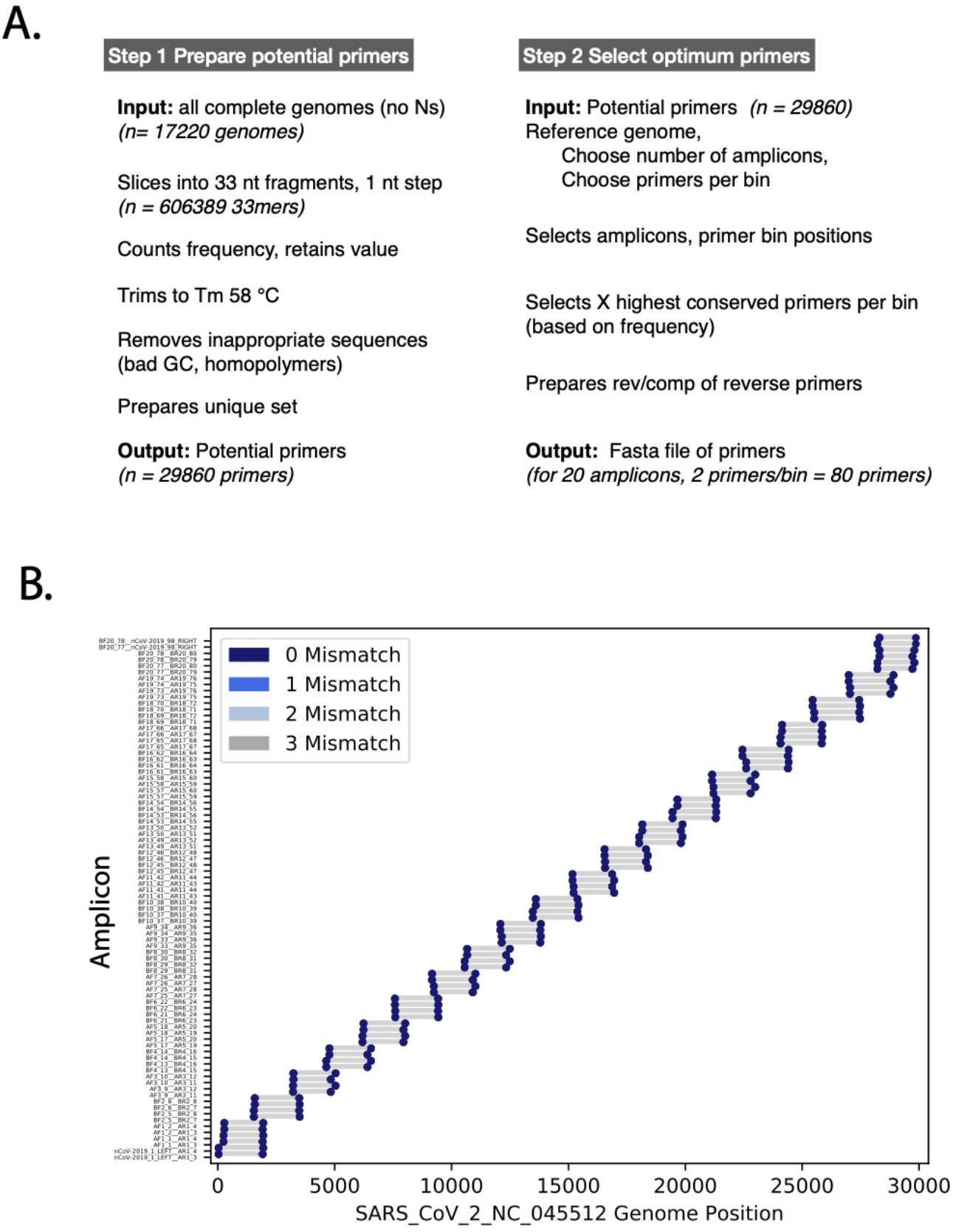
Primer design and amplicon layout. Panel A. The two main steps involved in primers generation and selection are shown. Panel B. The layout of the 20 amplicons across the SARS-CoV-2 genome is shown in lower panel. The blue markers indicate target positions in the SARS-CoV-2 genome (NC_045512 used here), the grey bars indicate the resulting amplicon.

In the second step we defined forward and reverse primer target regions (bins) for the amplicons. For SARS-CoV-2, we selected 20 amplicons with an overlap of 300nt, regularly spaced across the SARS-CoV-2 genome sequence (Figure 3B). We then selected the top conserved primer sequences (the highest frequency primers) mapping in the 5’ or 3’ 185nt of each amplicon. For security, the two highest frequency primers per bin were selected for the SARS-CoV-2 sequence, this provided some insurance against primer failure either due to target evolution or unexpected secondary structure. The binning and primer target locations for the final set of primers are shown in Figure 2 and the final calculated amplicon lengths were 1495-2093nt.

The reverse transcription, PCR amplification and library protocols were modified to accommodate the new primers. Important changes to note are the following. Reverse transcription was performed using the reverse primers and reverse transcription at 42°C. The PCR cycling conditions (using Phusion enzyme) were adjusted for the new T_m_s and an increased elongation time required for the longer PCR products. Finally, the library purification steps were adjusted to recover longer PCR and library products. A detailed step-by-step protocol is provided in the supplementary material.

## Testing the performance of primers to sequence SARS-CoV-2 using MinION

We tested the Entebbe primers performance for sequencing SARS-CoV-2 from nucleic acid extracted from positive samples. The amplicon sizes and genome coverage are summarised in Figure 4. In particular, panel A and B (Figure 4) illustrate the amplicon products after the reverse transcription and PCR amplification with the expected sizes of 1400-2000 bp. These amplicons were then pooled and used for library preparation using the MinION sequencing kits SQK-LSK109. Final libraries were quantified and sequenced using a MinION Flow Cell (R9.4.1). The resulting read data, after quality and primer and adapter trimming, were then mapped to the SARS-CoV-2 Wuhan1 reference genome NC_045512 (Figure 4, panels C and D) to document sequence coverage across the genome. The twenty individual amplicons are detected in the coverage pattern with small peaks appearing where amplicons overlap. The coverage is consistent across the genome with no missing amplicons and the data were readily assembled into good coverage full genomes. Initial experiments showed that amplicons 2 (spanning nt 2,400) and 16 (spanning nt 23,500) had reduced yields (Figure 4, panel C). The primer mixes were subsequently adjusted to increase concentrations of amplicon 1 and 16 primers for reverse transcription and PCR (see detailed protocol in Supplementary materials), this improved yields relative to the other amplicons (Figure 4 panel D).

**Figure 4.**
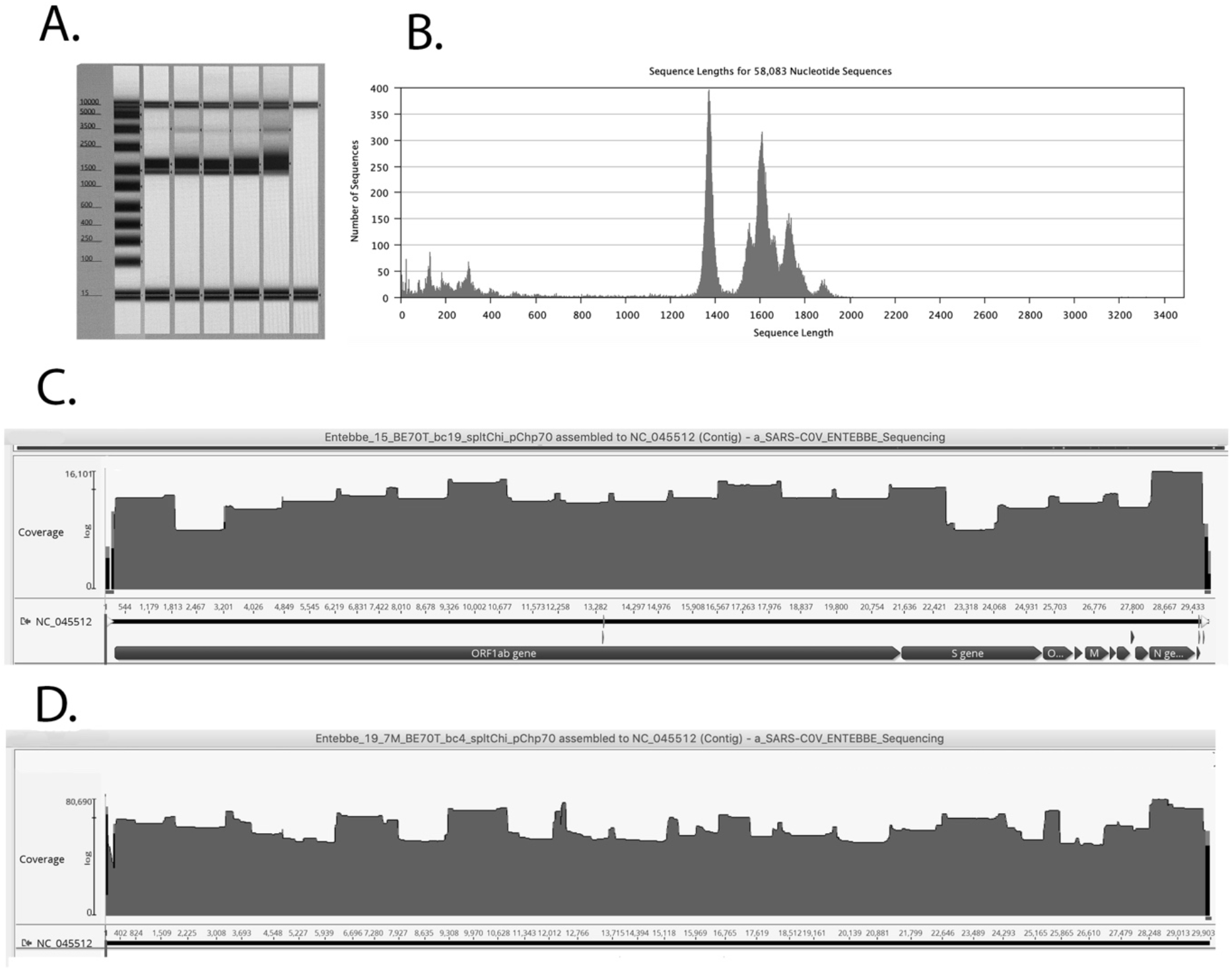
Testing the primer performance. Panel A, PCR product size after pooling of reaction A and B. Expected sizes of amplicons are from 1500 bp to 2093 bp before primer trimming. Panel B, MinION reads after quality control, primer, adapter trimming. Panel C, Reads mapped to SARS-CoV-2 reference genome, before amplicon 2 and 16 primer boosting. Panel D, Reads mapped to SARS-CoV-2 reference genome, after amplicon 2 and 16 primer boosting.

A set of SARS-CoV-2 clinical samples were tested with the Entebbe primers/protocol. Respiratory swab samples from 111 PCR confirmed cases of SARS-CoV-2 infection were processed for reverse transcription/PCR using the Entebbe primers and protocol as described in Supplementary Materials. If sufficient amplicon DNA was generated after PCR, MinION libraries were prepared, samples were sequenced on the MinION flowcells and the resulting data were assembled into genomes. Figure 5A shows the results of this validation test. Complete genomes (fraction genome= 1) were obtained from samples up to Ct 35, 19 samples failed to yield sufficient amplicon DNA after PCR stage (figure 5A, red markers) and 5 samples yielded genomes with gaps > 842 nt. The PCR failures and the gapped genomes were not strictly associated with higher Cts and their distribution pattern is similar to the overall Ct distribution pattern across the set of samples (Figure 5 panel B) suggesting other factors such as sample quality, extraction method, storage, might be more critical than Ct in determining sequencing success, at least in samples within the Ct range tested (up to Ct 37).

**Figure 5.**
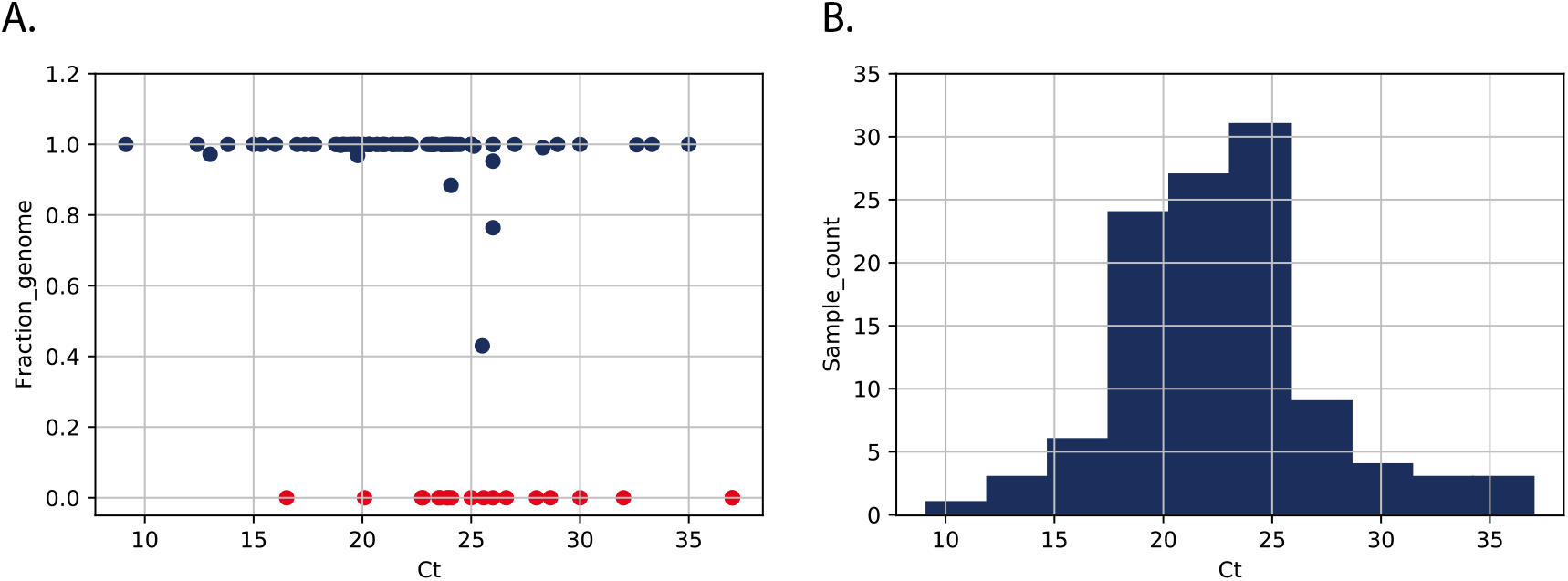
Validation of Entebbe primers. **Panel A** plots the genome yield (fraction of complete genome) as a function of sample Ct. Fraction genome was calculated by number of nonN nucleotides/29303 (the length, in nt, of NC_045512 reference genome). Each marker represents a sample, red markers indicate 19 samples that failed to yield sufficient DNA for library, 93 that proceeded to library preparation and sequencing (dark blue markers). **Panel B** is a histogram of the distribution of the 118 sample Cts.

## Conclusions

Given the urgency of controlling the SARS-CoV-2 pandemic and the importance of having good quality SARS-CoV-2 genomes, we are providing these alternative primers (the Entebbe primers) with detailed step-by-step laboratory protocols to the community with the hope that they benefit from the new design. The costs and efforts of sequencing SARS-CoV-2 in the large case numbers that are currently being seen are substantial and if these new primers result in a higher proportion of gap-less genomes, this will provide added value and will increase the utility of the resulting data.

## Acknowledgements

We thank all global SARS-CoV-2 sequencing groups for their open and rapid sharing of sequence data and GISAID for providing an effective platform for making these data available. We are grateful to the Oxford Nanopore Technologies and the ARTIC Network for their support with protocols and analysis software. We thank the Central Public Health Laboratory, Uganda especially Isaac Ssewanyana, Patrick Semanda, Susan N. Nabadda for their support with SARS-CoV-2 samples.

We acknowledge the support of the Uganda Ministry of Health and its COVID-19 Scientific Advisory Committee, the National COVID-19 Task Force, and the staff of the Emerging and Remerging Infections Department of the Uganda Virus Research Institute, and the US Centers for Disease Control and Prevention. The SARS-CoV-2 diagnostic and sequencing award is jointly funded by the UK MRC and the UK DFID under the MRC–DFID Concordat agreement (grant agreement no. NC_PC_19060) and is also part of the European and Developing Countries Clinical Trials Partnership 2 program supported by the European Union. The diagnostics were also supported by the UK Research and Innovation/MRC, the Global Fund, the Government of Uganda, the Islamic Development Bank, the World Health Organization, GAVI, the US Centers for Disease Control and Prevention, and the Jack Ma Foundation, among others. M.V.T.P. was supported by a Marie Sklodowska-Curie Individual Fellowship, funded by European Union’s Horizon 2020 research and innovation programme (grant agreement no. 799417). The Uganda Medical Informatics Centre high performance computer was supported by the UK MRC (grant no. MC_EX_MR/L016273/1) to P.K. The study is supported by a Wellcome Epidemic Preparedness–Coronavirus grant, jointly funded by the Wellcome Trust and UK DFID (grant agreement no. 220977/Z/20/Z) awarded to M.C. This study was approved by the UVRI Research and Ethics Committee (approval no. 00001354, study reference no. GC/127/20/04/771).

---

**Supplementary Materials: 1. Supplementary Material Figure 1, 2. Detailed protocol, 3. Primer table.**

**Supplementary Material Figure 1.**
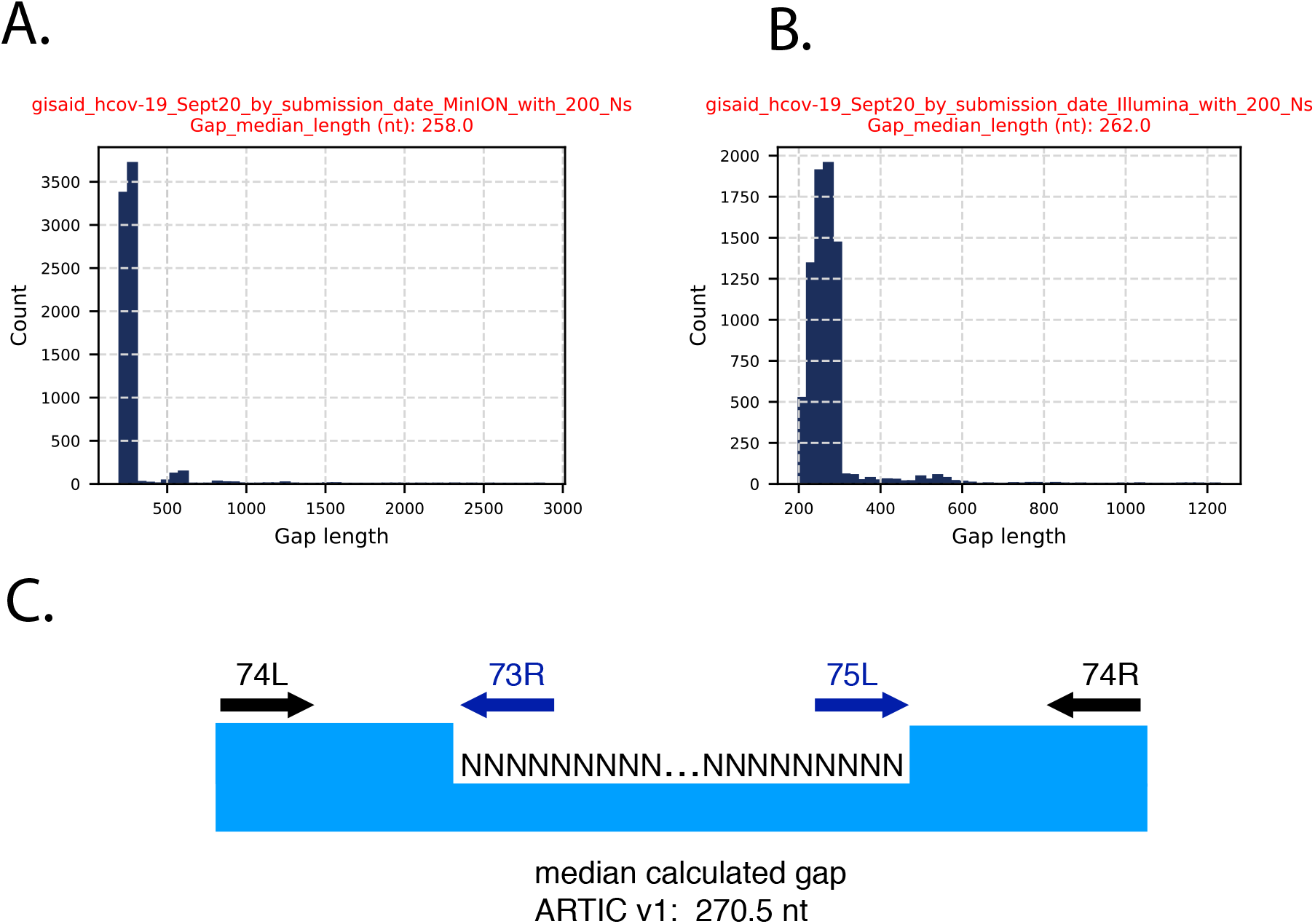
As described in Figure 1, SARS-CoV-2 genomes deposited in September 2020 (n= 38,228) were retrieved from GISAID, sorted by sequencing platform and the presence of 200N motifs was monitored. Panel A presents a histogram of gap lengths present in the first 3000 MinION-generated genome sequences that contained at least one 200N motif. The median length for all gaps was 258 nt. Panel B present the same analysis for Illumina-generated genome sequences. Panel C presents a typical gap. Because of overlapping amplicons, if a single amplicon is missing (amplicon 74 for example) from the sequencing library, the resulting gap in coverage would not be the complete amplicon 74 but would span from the 3’ end of the adjacent amplicon 73 (after primer and quality trimming) to the 5’ end of adjacent amplicon 75 (after primer and quality trimming). The calculated gaps generated by such amplicon loss have a median length of 270.5 nt, which is close to the the observed median gap length in the MinION data (258 nt) or Illumina data (262 nt) from September 2020.

## 2. Detailed protocol

We acknowledge the Oxford Nanopore Technologies and the ARTIC Network for their extensive protocols which provided important background details for the method presented here (19)(20). Potential users are strongly advised to study the original protocols carefully. The protocol presented here assumes the users has all the ARTIC and ONT background experience.

### Detailed protocol for reverse transcription, PCR and sequencing library preparation

#### Practical comments

Primers were ordered and delivered dry and were dissolved at 100 *μM* (= 100 pmol/*μ*l) in PCR quality water. Dry shipped primers are reported to have better stability at room temperature (21). Ideal the primers were allowed to dissolve in PCR grade water for at least 3 hours at room temperature or overnight at 4 °C before diluting for use (see below).

##### Prepare the following primer mixes

**RPM_A**: 21 primers. 3 *μ*l of 100 *μM* stock of the following Reverse primers plus PCR grade water to 200 *μ*l (= 300 pmol x 21 = 6300 pmol/200 *μ*l = 32 pmol/*μ*l. 2 *μ*l per 20 *μ*l RT reaction = 3.2 pmol/*μ*l = 3.2 *μM* concentration in reaction).

**Primers in RPM_A**: nCoV-2019_1_LEFT, AR1_3, AR1_4, AR3_11, AR3_12, AR5_19, AR5_20, AR7_27, AR7_28, AR9_35, AR9_36, AR11_43, AR11_44, AR13_51, AR13_52, AR15_59, AR15_60, AR17_67, AR17_68, AR19_75, AR19_76.

**RPM_B**: 21 primers. 3 *μ*l of 100 *μM* stock of the following Reverse primers plus PCR grade water to 200 *μ*l. (= 300 pmol x 21 = 6300 pmol/200 *μ*l = 32 pmol/*μ*l. 2 *μ*l per 20 *μ*l RT reaction = 3.2 pmol/*μ*l = 3.2 *μM* concentration in reaction). **Primer Boost:** For Primers BR2_7, BR2_8, BR16_63, BR16_64 use 9 *μ*l of 100 *μM* stock.

**Primers in RPM_B:** BR2_7, BR2_8, BR4_15, BR4_16, BR6_23, BR6_24, BR8_31, BR8_32, BR10_39, BR10_40, BR12_47, BR12_48, BR14_55, BR14_56, BR16_63, BR16_64, BR18_71, BR18_72, BR20_79, BR20_80, nCoV-2019_98_RIGHT.

**PCR primer mix A (PPM_A):**1.5 *μ*l of each primer (forward and reverse primer) plus 439 *μ*l PCR grade water. = 41 primers (100 pmol/*μ*l) = 4100 pmol in 500 *μ*l water = 8.2 pmol/*μ*l in the PPM. When 2 *μ*l PPM used per 25 *μ*l PCR reaction = 0.66 pmol/*μ*l in reaction (= 660 nM).

**Primers in PPM_A:** nCoV-2019_1_LEFT, AF1_1, AF1_2, AR1_3, AR1_4, AF3_9, AF3_10, AR3_11, AR3_12, AF5_17, AF5_18, AR5_19, AR5_20, AF7_25, AF7_26, AR7_27, AR7_28, AF9_33, AF9_34, AR9_35, AR9_36, AF11_41, AF11_42, AR11_43, AR11_44, AF13_49, AF13_50, AR13_51, AR13_52, AF15_57, AF15_58, AR15_59, AR15_60, AF17_65, AF17_66, AR17_67, AR17_68, AF19_73, AF19_74, AR19_75, AR19_76.

**PCR primer mix B (PPM_B):**1.5 *μ*l of each primer (forward and reverse primer) plus 439 *μ*l PCR grade water = 41 primers (100 pmol/*μ*l) = 4100 pmol in 500 *μ*l water = 8.2 pmol/*μ*l in the PPM. When 2 *μ*l PPM used per 25 *μ*l PCR reaction = 0.66 pmol/*μ*l in reaction (= 660 nM). **Primer Boost:** for primers BF2_5, BF2_6, BR2_7, BR2_8, BF16_62, BF16_61, BR16_63, BR16_64, use 4.5 *μ*l of 100 *μM* stock

**Primers in PPM_B:** BF2_5, BF2_6, BR2_7, BR2_8, BF4_13, BF4_14, BR4_15, BR4_16, BF6_21, BF6_22, BR6_23, BR6_24, BF8_29, BF8_30, BR8_31, BR8_32, BF10_37, BF10_38, BR10_39, BR10_40, BF12_45, BF12_46, BR12_47, BR12_48, BF14_53, BF14_54, BR14_55, BR14_56, BF16_61, BF16_62, BR16_63, BR16_64, BF18_69, BF18_70, BR18_71, BR18_72, BF20_77, BF20_78, BR20_79, BR20_80, nCoV-2019_98_RIGHT.

###### 1. Set up Reverse Transcription reactions (two reactions per sample)

Mix the following components in a 0.2mL 8-strip tube, 1 RT_A and 1 RT-B reaction per sample.

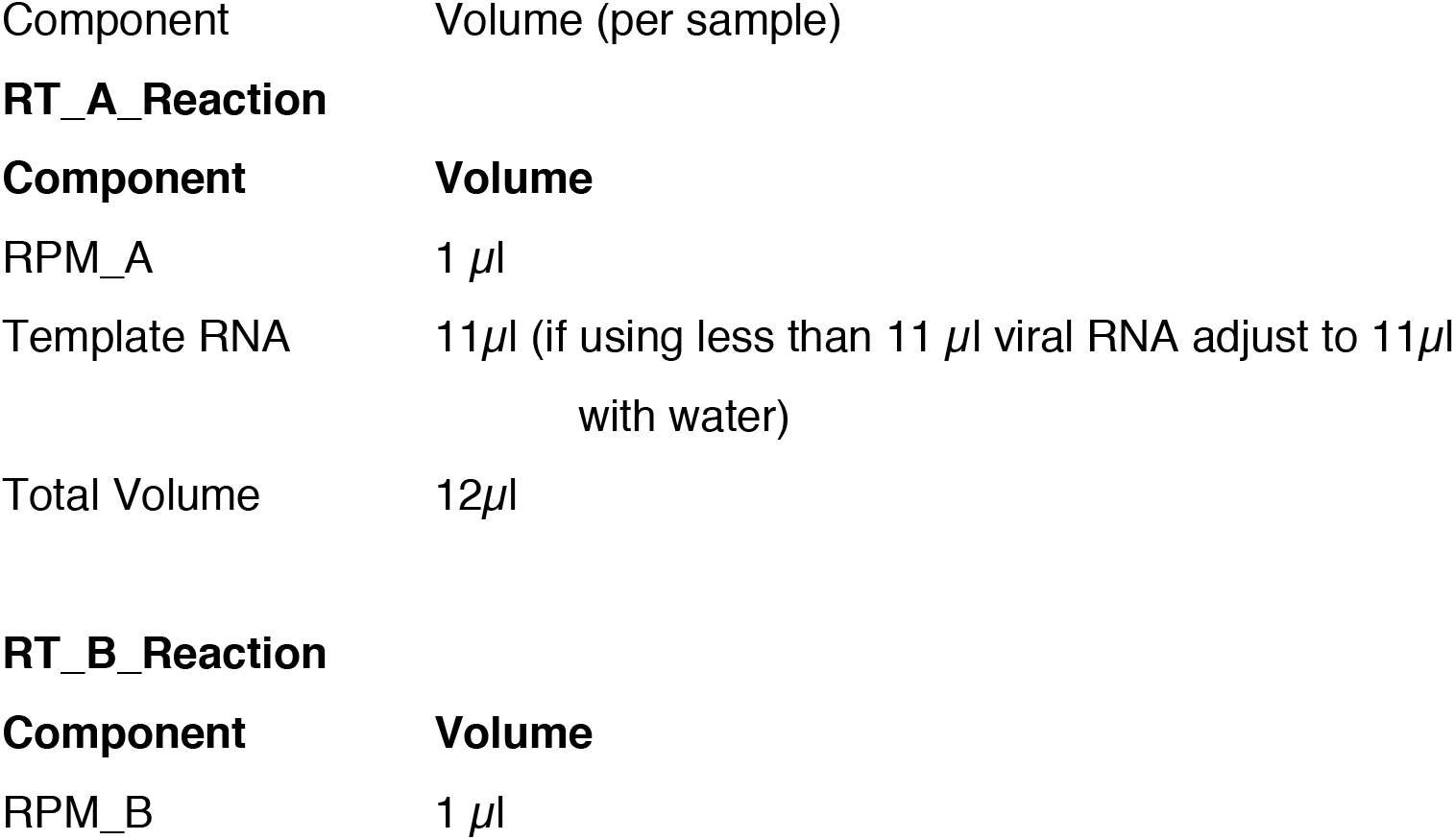
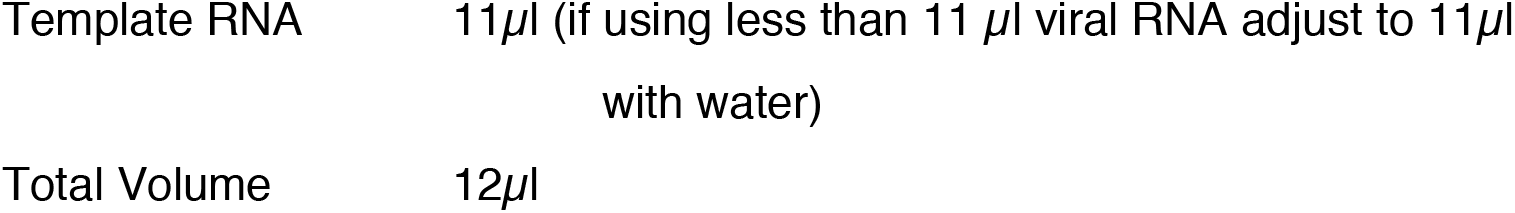

Heat at 65 °C for 5:00 mins, and quickly chill in ice/water bath for at least 1:00 min.

Viral RNA input from a clinical sample should be between Ct 18-35. If Ct is between 12-15, then dilute the sample 100-fold in water, if between 15-18 then dilute 10-fold in water. This will reduce the likelihood of PCR-inhibition.

###### Reverse Transcription

Add the following to the 12 *μ*l annealed template RNA sample:

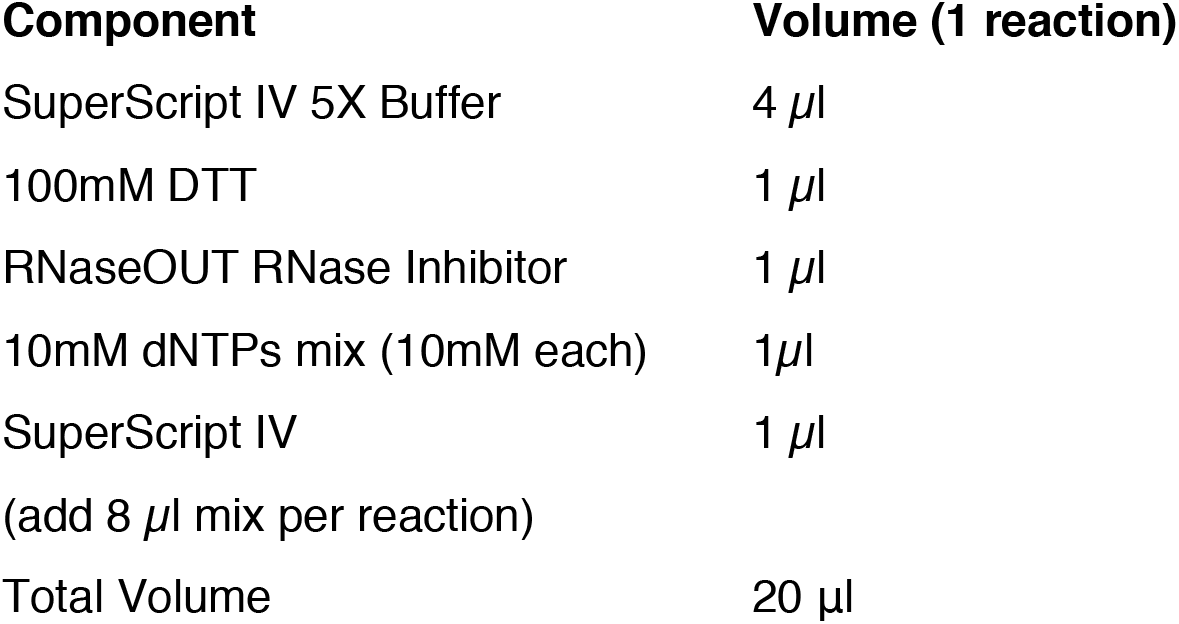

| 430 | Component | Volume (1 reaction) |
| --- | --- | --- |
| 434 | 10mM dNTPs mix (10mM each) | 1μl |
| 437 | Total Volume | 20 μl |

###### 2. Incubate the RT reaction as follows

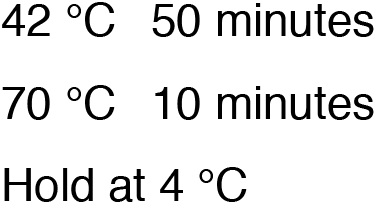

###### 3. Set up Amplicon PCR

In the Mastermix hood set up the multiplex PCR reactions as follows in 0.2mL 8-strip PCR tubes. Prepare one PPM_A and one PPM_B mix per sample.

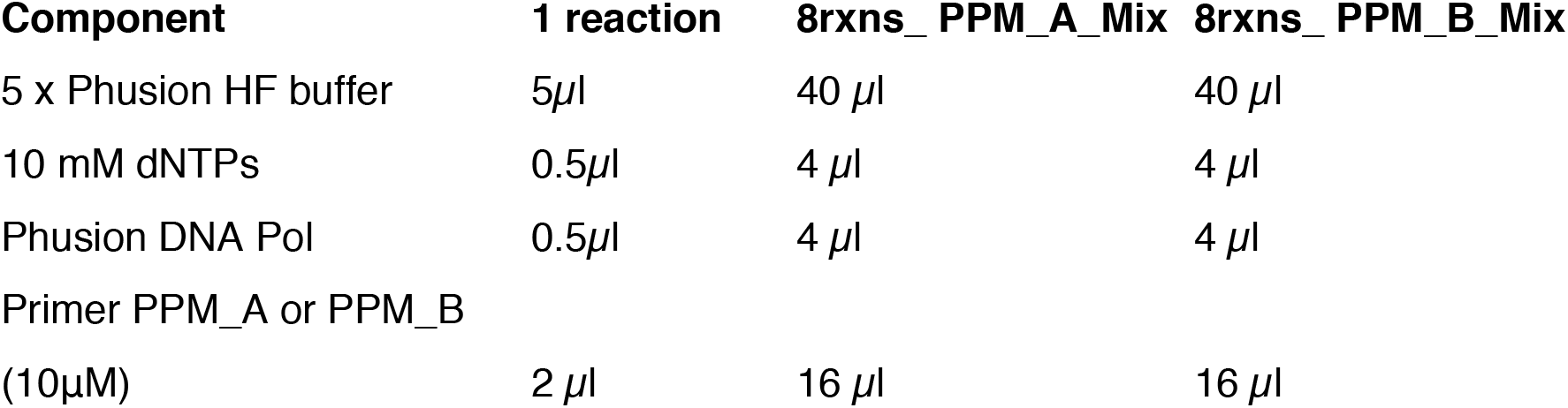
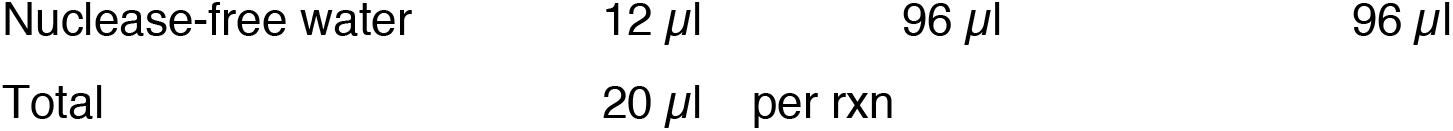

Aliquot 20 *μ*l per tube PPM_A_Mix or PPM_B_Mix

In the extraction and sample addition cabinet add **5μl cDNA** to each tube and mix well by pipetting.

###### 4. Run PCR

| 462 **Step** | **Cycles** | **Temp** | **Time** |
| --- | --- | --- | --- |
| 463 Heat Activation | 1 | 98 °C | 30sec |
| 464 Denaturation | 35 | 98 °C | 15sec |
| 465 Annealing | 35 | 63 °C | 30sec |
| 466 Extension | 35 | 72 °C | 3 min |
| 467 Polish | 1 | 72 °C | 10 min |
| 468 Hold |  | 4 °C | infinite |

In post-PCR cabinet: After PCR, for each sample, pool the two reactions in a 1.5 ml microcentrifuge tube.

###### 5. Clean up using AMPure XP beads, Agencourt

Add an equal volume (1:1) of AMPure XP beads to the sample tube and mix gently by either flicking or pipetting. E.g. add 50 μl AMPure XP bead suspension to a 50 μl reaction.

Pulse centrifuge to collect all liquid at the bottom of the tube.

Incubate for 5 min at room temperature.

Place on magnetic rack and for 2 min

Carefully remove and discard the supernatant, being careful not to touch the bead pellet.

**Ethanol wash 1:** Add 200μl of room-temperature 70%volume ethanol to the pellet. Carefully remove and discard ethanol, being careful not to touch the bead pellet.

###### Repeat Ethanol wash

Pulse centrifuge to collect all liquid at the bottom of the tube and carefully remove as much residual ethanol as possible using a P10 pipette.

Open lid, allow to dry briefly (2 min)

Resuspend pellet in 30μl Elution Buffer (EB, QIAGEN)

Mix gently by flicking,

Incubate 5 min

Place on magnet and transfer sample (supernatant) to a clean 1.5mL Eppendorf tube ensuring no beads are transferred into this tube.

###### 6. uantitate

(A Tapestation 4200 analyzer with the D5000 assay kit as well as the Qubit fluorometer were used.)

###### 7. Add Barcodes (Steps 7 onward derived from the Oxford Nanopore Technologies protocol (19))

Set up the following reaction for each sample:

| 505 **Component** | **Volume** |
| --- | --- |
| 506 DNA amplicons | 5 µl* |
| 507 Nuclease-free water | 7.5 µl |
| 508 Ultra II End Prep Reaction Buffer | 1.75 µl |
| 509 Ultra II End Prep Enzyme Mix | 0.75 µl |
| 510 Total Volume | 15 µl |

\*Volumes usually varies based on the DNA concentration. For the 1500bp amplicons and
pool size of appx 8 we have input 50ng and for a pool size of 12 we have input 20ng

Incubate at room temp (20°C) for 20 min

Incubate at 65 °C for 5:00 min

Incubate on ice for 1 min

E7645S/L NEBNext Quick Ligation Module

| 520 **Component** | **Volume** |
| --- | --- |
| 521 End repaired sample | 15µl |
| 522 Barcode vol | 2.5 µl |
| 523 Ultra II Ligation Master Mix | 17.5 µl |
| 524 Ligation Enhancer | 0.5 µl |
| 525 Water | 1.5 µl |
| 526 final volume | 35.5 µl |

Incubate in a thermocycler for 15 minutes at 20°C with heated lid set to 30°C (or off).

Incubate at 70 °C for 10 min

Incubate on ice for 1 min

The 70°C incubation is to inactivate the DNA ligase to prevent barcode cross-ligation when reactions are pooled in the next step.

Pool all barcoded fragments together into a new 1.5 ml Eppendorf tube.

###### 8. Clean up on AMPure XP beads, Agencourt

Add an equal volume (1:1) of SPRI beads to the sample tube and mix gently by either flicking or pipetting. E.g. add 50 μl SPRI beads to a 50 μl reaction.

Pulse centrifuge to collect all liquid at the bottom of the tube.

Incubate for 5 min at room temperature.

Place on magnetic rack and for 5 min

Carefully remove and discard the supernatant, being careful not to touch the bead pellet.

**Ethanol wash 1:** Add 200μl of room-temperature 70%volume ethanol to the pellet. Carefully remove and discard ethanol, being careful not to touch the bead pellet.

###### Repeat Ethanol wash

Pulse centrifuge to collect all liquid at the bottom of the tube and carefully remove as much residual ethanol as possible using a P10 pipette.

Open lid, allow to dry briefly (2 min)

Resuspend pellet in 30μl Elution Buffer (EB, QIAGEN)

Mix gently by flicking

Incubate 5 min

Place on magnet and transfer sample (supernatant) to a clean 1.5mL Eppendorf tube ensuring no beads are transferred into this tube.

###### 9. Quantitate as above

###### 10. Add AMII adapters

AMII adapters ligation reaction:

| 571 | Component | Volume |
| --- | --- | --- |
| 572 | Barcoded amplicon pools | 30 μl |
| 573 | NEBNext (E60562) |  |
| 574 | Quick Ligation Reaction Buffer (5X) | 10 μl |
| 575 | AMII adapter mix | 5 μl |
| 576 | Quick T4 DNA Ligase | 5 μl |
| 577 | Total | 50 μl |

The input of barcoded amplicon pools will depend on the number of barcoded pools and should be between 40 ng (8 barcodes) and 160 ng (24 barcodes).

Incubate at room temperature for 15:00 min

###### 11. Clean up on AMPure XP beads, Agencourt

**USE SFB instead of Ethanol for two washes ***CRUCIAL,** Ethanol strips off motor protein from adapter. You don’t want to do this!

Clean up on AMPure XP beads, Agencourt

Add an equal volume (1:1) of SPRI beads to the sample tube and mix gently by either flicking or pipetting. E.g. add 50 μl SPRI beads to a 50 μl reaction.

Pulse centrifuge to collect all liquid at the bottom of the tube.

Incubate for 5 min at room temperature.

Place on magnetic rack and for 2 min

Carefully remove and discard the supernatant, being careful not to touch the bead pellet.

**SFB washes:** Add 200μl of room-temperature **SFB** to the pellet.

resuspend

collect on magnet

Carefully remove and discard **SFB** being careful not to touch the bead pellet.

###### Repeat SFB wash

Pulse centrifuge to collect all liquid at the bottom of the tube and carefully remove as much residual **SFB** as possible using a P10 pipette.

###### Resuspend pellet in 15 *μ*l Elution Buffer (EB, QIAGEN)

Mix gently by flicking,

###### Incubate 5 min

Place on magnet and transfer sample (supernatant) to a clean 1.5mL Eppendorf tube ensuring no beads are transferred into this tube.

###### 12. Quantitate

###### 13. Proceed to MinION sequencing

**Supplementary material Table 1.**
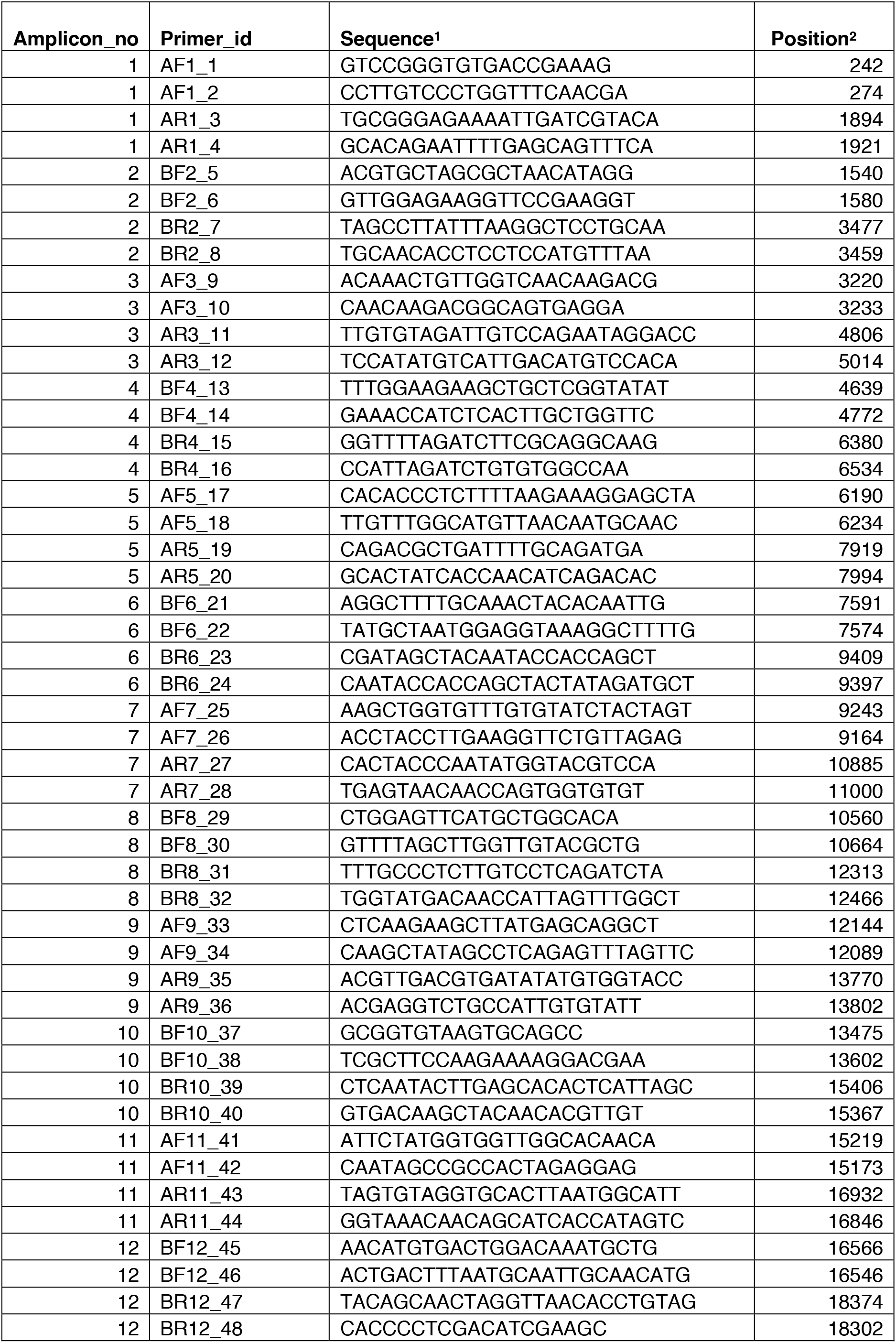
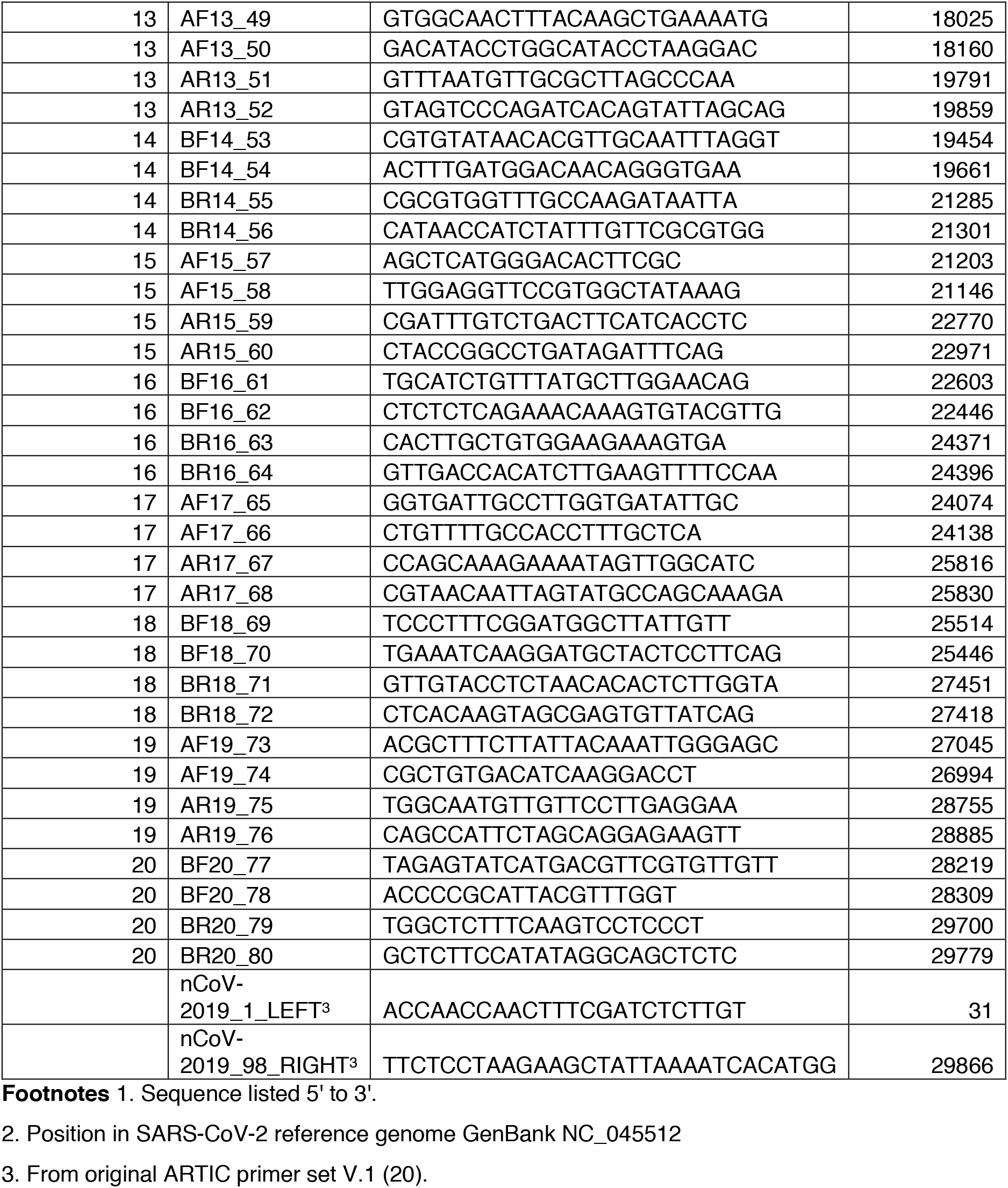
Entebbe Primers

## Notes

### Competing Interest Statement

The authors have declared no competing interest.

### Summary of Updates

Manuscript revised to address peer reviewers's comments. Revisions included improved resolution of graphics in Figure 1 and 2, additional Figure 5 with testing of primers on a larger set of clinical samples, correction of typos and grammatical errors.

